# Structural basis of multitasking by the apicoplast DNA polymerase from *Plasmodium falciparum*

**DOI:** 10.1101/2025.04.09.647933

**Authors:** Anamika Kumari, Theodora Enache, Timothy D. Craggs, Janice D. Pata, Indrajit Lahiri

## Abstract

*Plasmodium falciparum* is a unicellular eukaryotic pathogen responsible for the majority of malaria-related fatalities. *Plasmodium* belongs to the phylum Apicomplexa and like most members of this phylum, contains a non-photosynthetic plastid called the apicoplast. The apicoplast has its own genome, which is replicated by a dedicated apicoplast replisome. Unlike other cellular replisomes, the apicoplast replisome uses a single DNA polymerase (apPol) for copying the apicoplast DNA. Being the only DNA polymerase in the apicoplast, apPol is expected to multitask, catalysing both replicative and lesion bypass synthesis. Replicative synthesis typically relies on a restrictive active site for high accuracy while lesion bypass requires an open active site. This raises the question how does apPol combine the structural features of multiple DNA polymerases in a single protein. Using single particle electron cryomicroscopy (cryoEM), we have solved the structures of apPol bound to its DNA and nucleotide substrates in five pre-chemistry conformational states, allowing us to describe the events leading up to nucleotide incorporation and answer how apPol incorporates features of multiple polymerases. We found that, unlike most replicative polymerases, apPol can accommodate a nascent base pair with the fingers in an open configuration, which might facilitate the lesion bypass activity. In the fingers open state we identified a nascent base pair checkpoint that can preferentially select a Watson-Crick base pair, an essential requirement for replicative synthesis. Taken together these structural features explain how apPol may balance replicative and lesion bypass synthesis.

## INTRODUCTION

*Plasmodium* is a parasitic protozoan responsible for causing malaria. This parasite belongs to the phylum Apicomplexa and like most members of this phylum, *Plasmodium* contains an essential, non-photosynthetic plastid called the apicoplast ^1^. *Plasmodium* apicoplast has its own circular 35kb A/T-rich genome coding for a chaperone as well as proteins and RNA involved in apicoplast gene expression ^2^. Apicoplast DNA is copied by a dedicated replication machinery called the apicoplast replisome. Compared to other cellular replisomes, the apicoplast replisome functions with very few proteins. For instance, all cellular replisomes use replicative polymerases to accurately copy undamaged DNA and specialised translesion synthesis (TLS) and repair polymerases to bypass damaged DNA and perform DNA repair respectively. Apicoplast replication, however, is carried out by a single DNA polymerase called apPol, indicating that this polymerase can perform the roles of multiple DNA polymerases within a single enzyme ^3–5^.

Replicative polymerases are fast and accurate enzymes and copy the bulk of the DNA. A major determinant of accurate synthesis is a restrictive active site favoring Watson-Crick base pairs ^6–8^. However, the accuracy comes at a cost, as replicative polymerases cannot efficiently copy damaged DNA or lesions ^9^. On encountering a lesion, replicative polymerases might stall, in turn stalling the replisome. If left unresolved, stalled replisomes can lead to replication fork collapse and eventual cell death. To rescue this situation one of the strategies used by cellular replisomes is to employ TLS polymerases to bypass the lesion, thus preventing replication stalling ^10^. The specialised lesion bypass functionality of TLS polymerases relies on a more open, promiscuous active site and comes at the cost of reduced accuracy ^9,11,12^. ApPol has the rare ability of performing both replicative and lesion bypass synthesis and it is a puzzle how this enzyme combines the differing active site requirements for these two activities.

The structures of most DNA polymerases solved to date resemble a right hand with the enzyme divided into the thumb, palm and fingers subdomains ^13,14^ (Figure 1).

**Figure 1:**
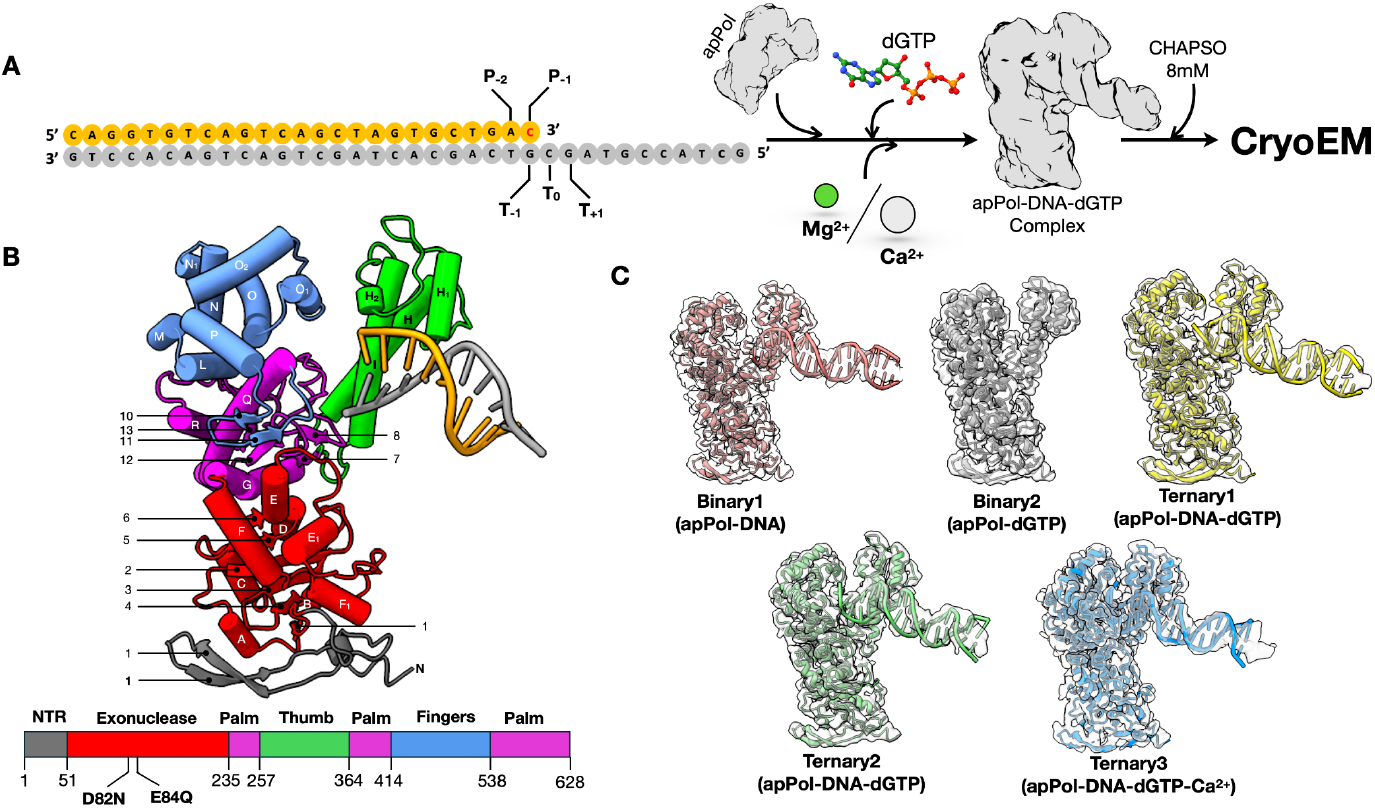
CryoEM of apPol pre-chemistry complexes. **(A)** apPol complex assembly scheme. A synthetic primer/template DNA (primer: orange; template: grey) was incubated with apPol and dGTP in the presence of either Mg^2+^ or Ca^2+^ to form apPol pre-chemistry complex. 8mM CHAPSO was added just before cryoEM grid preparation. The templating base (T_0_) and the ones immediately 5’ and 3’ to it (T_+1_ and T_-1_ respectively) along with the two bases at the 3’ end of the primer strand (P_-2_ and P_-1_) are highlighted. The P_-1_ base (dC; red) is dideoxy terminated when Mg^2+^ is used as the divalent metal. **(B)** Atomic model (tube and arrow representation) of the 3.2Å consensus map from dataset1 along with the domain organisation of apPol. The domains are coloured as follows (N-terminal region (NTR): grey; 3’-5’ proofreading exonuclease: red; palm: magenta; thumb: green and fingers: blue). The exonuclease active site residues (D82 and E84) that have been mutated to N and Q respectively in all the apPol constructs used in this study are highlighted. The secondary structural elements are labelled based on structure alignment with Klenow fragment (PDB id: 1KLN)^14^. **(C)** Atomic models of the two binary (apPol-DNA (binary1): salmon pink; apPol-dGTP (binary2): grey) and three ternary complexes of apPol-DNA-dGTP (ternary1, ternary2 and ternary3: yellow, green and blue respectively) fitted to their respective cryoEM maps (transparent grey).

Typically the thumb interacts with the DNA duplex, stabilizing the DNA on the polymerase. The catalytic aspartic acid residues involved in the two metal ion mechanism of nucleotide addition ^15^ are located in the palm domain while the fingers is involved in ensuring accuracy of the DNA synthesis. The replicative polymerases possess an additional 3’ to 5’ exonuclease activity, which resides in the proofreading exonuclease domain of these enzymes.

Most DNA polymerases, including apPol, follow the same overall catalytic cycle for DNA synthesis ^5,16–19^ (Figure S1A). The pre-chemistry steps involve binding of DNA and nucleotide substrate (Figures S1A, steps 1 and 2) followed by the transition to a catalytically competent pre-chemistry ternary complex (Figures S1A, step 3). In most replicative polymerases, this transition involves a large-scale conformational change in the fingers domain termed fingers closure. Fingers closure leads to the formation of a restrictive active site, favouring a Watson-Crick base pairing between the templating base (T_0_) and the incoming dNTP ^7,20^. Fingers closure is followed by the chemical step of nucleotide addition (Figure S1A, step 4) using the two-metal-ion or three-metal-ion mechanism ^15,21,22^. The post-chemistry steps involve the release of the inorganic pyrophosphate byproduct (PP_i_) followed by translocation along the DNA (Figure S1A, steps 4 and 5 respectively).

Apicoplast DNA undergoes reactive oxygen species mediated oxidative damage and, since apPol is the only polymerase in the apicoplast, this enzyme is anticipated to perform both replicative and lesion bypass DNA synthesis ^4^. In fact, kinetic analysis shows that apPol can bypass oxidative lesions including 8-oxo-7,8-dihydro-2’-deoxyguanosine (8oxodG), 5,6-dihydroxy-5,6-dihydro-2’-deoxythymine (thymine glycol) and 2-oxo-1,2-dihydro-2’-deoxyadenine (2oxodA) ^4,5^. In TLS polymerases, the fingers closing is less pronounced or absent altogether ^11,12,23^, leading to a wider active site that can accommodate lesions and bypass them. However, the wide active sites allows TLS polymerases to accommodate non Watson-Crick base pairs while copying undamaged DNA, thus making these enzymes error prone. Since the structural requirements for replicative and lesion bypass synthesis are very different it raises the question how does apPol perform both replicative and lesion bypass synthesis?

ApPol belongs to the A-family of DNA polymerases but is only distantly related to the well studied A-family polymerases including bacterial Pol I, T7 DNA polymerase and mitochondrial DNA polymerase gamma ^24^. Instead, apPol belongs to a poorly studied clade of A-family DNA polymerase and members of this clade are found in all domains of life^25^. Recent structures of apo, binary and pre-chemistry ternary complexes of apPol show how the enzyme interacts with its substrates ^26,27^. However, these structures do not shed light on how apPol might accommodate both replicative and translesion synthesis.

We have used electron cryomicroscopy (cryoEM) to solve a series of structures of apPol in complex with a primer/template DNA and the correct incoming dNTP (Figure 1). These structures elucidate the conformational changes of apPol critical for the pre-chemistry steps. We show that, unlike other A-family polymerases, the pairing between the templating base (T_0_) and incoming nucleotide occurs with the fingers domain of apPol in the open conformation. Moreover, we propose a molecular checkpoint specific to apPol that might allow this polymerase to perform TLS without sacrificing the accuracy needed for replicative synthesis.

## RESULTS

We assembled two pre-chemistry ternary complexes of apPol (with an inactive exonuclease domain) bound to a primer/template DNA and the corresponding incoming nucleotide (dGTP) (Figures 1A and B). To prevent the chemical step of bond formation (Figure S1A step 3) we either used a 3’ dideoxy-terminated primer strand with Mg^2+^ as the divalent cation (dataset1) or a non dideoxy-terminated primer strand with Ca^2+^ as the cation (dataset2). In both datasets 8 mM CHAPSO was used to achieve a uniform particle distribution over holey carbon grids (Figure 1A). We performed a multiple nucleotide incorporation assay to ascertain that apPol retained its activity in the presence of CHAPSO (Figure S1B).

We solved the structures of these complexes using cryoEM. For dataset1 the consensus structure was determined to an overall resolution of 3.2Å (Figure S2). The apPol density was well-resolved and we could trace the entire polypeptide backbone (Figure 1B) and identify densities for several amino acid residues. Overall, the apPol structure was similar to the previously reported structure of the apo enzyme ^26^. The DNA density was fragmented and there was no clear density for the incoming nucleotide (Figures 1B and S2). We further classified the particles contributing to the consensus map and could resolve them into four structures (global resolution ranging from 3.8Å to 4.1Å) corresponding to apPol-DNA, apPol-dGTP, and two conformational states of apPol-DNA-dGTP complexes (Figures 1C and S2). From dataset2 we solved an apPol-DNA-dGTP-Ca^2+^ ternary complex structure to a global resolution of 3.5Å (Figures 1C and S2). Here both the DNA and nucleotide substrates were clearly visible. By combining all these structures, we describe the events leading up to the chemical step of bond formation by apPol.

### apPol binds DNA and nucleotide independent of each other

The 3D classification of the particles contributing to the consensus structure of dataset1 resulted in two binary complexes, one had apPol bound to DNA (binary1) and the other had apPol bound to the incoming dGTP (binary2) (Figures S2 and 1C). In binary1 we could detect density for 20 out of the 25 base pairs of the DNA duplex (Figures 1C and S3A). Of the 20 base pairs, only 6 were contacted by apPol (Figure 2A).

**Figure 2:**
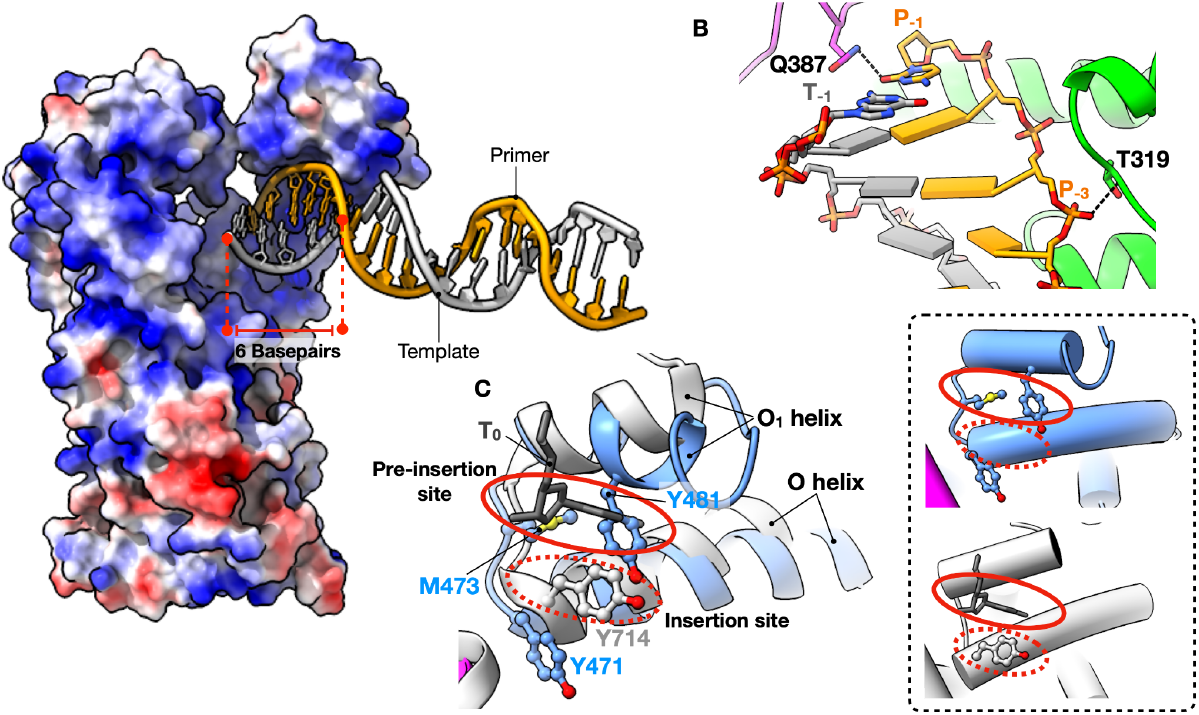
Binary1 complex of apPol. **(A)** 6 base pairs (stick representation; red dashed lines) of DNA interact with a positively charged groove (blue) of apPol (surface representation; coloured according to coulombic potential; red: negative charge to blue: positive charge). **(B)** Specific interactions of apPol with the DNA duplex. Colour coding is the same as Figure 1B. Potential hydrogen bonds are shown as black dashed lines. **(C)** Close up view of the pocket between the O and O1 helices of the fingers domain (blue) superimposed on the BSt Pol I binary structure (PDB id: 1L3S; white). The pre-insertion and insertion sites identified for BSt Pol I are highlighted with solid red and dashed red ovals respectively. Y481, M473 and Y471 of apPol and Y714 of BSt Pol I are shown in ball and stick representation. The T_0_ nucleotide of BSt Pol I is shown in dark grey. **Inset:** The pocket between O and O1 helices of apPol binary1 (top) and 1LS3 (bottom) shown individually. For simplicity, helices are shown as tubes.

In the sharpened map no clear density was visible for the template overhang including the T_0_ base. However, in the unsharpened map we could detect the density for the T_0_ base (Figure S3A) and that guided the register of the DNA duplex. Similar to other A family polymerases, the DNA is mainly contacted by the thumb domain of apPol. Helices H, H_1_ and H_2_ form a positively charged groove that accommodates the minor groove phosphate backbone of the DNA duplex (Figure 2A). However, consistent with a recent report ^27^, unlike other A family polymerases, the interactions between the DNA and the thumb is limited in apPol with a total buried surface area of only 607Å^2^. We detected two specific contacts, Q387 forms a hydrogen bond with the 3’ base (P_-1_) of the primer strand potentially determining the position of the DNA in binary1 and the backbone nitrogen of T319 interacts with the phosphate backbone of P_-3_ on the primer strand (Figure 2B). The template strand does not make any significant contact with apPol with a buried surface area of only 174Å^2^. We note that at the resolution range of the structures reported here (3.2Å to 4.1Å) it is not possible to reliably detect density for ordered water molecules and thus any potential water mediated interactions have not been analysed throughout this work.

The binary2 structure is similar to that of binary1 except that there is no clear DNA density (Figures 1C and S2). Instead, we can detect the density for the incoming dGTP near the O helix of the fingers domain (Figure S3C). The base moiety is stabilised by H-bond to Y486 of the O1 helix while the triphosphate forms hydrogen bonds with R390.

For most DNA polymerases, a transition from the apo to the catalytically poised pre-chemistry ternary complex is accompanied by the closure of the fingers domain ^28–32^, restricting the active site by forming a lid over the nascent base pair. For A-family polymerases, fingers closure is achieved by a large-scale rotation of the O and N helices towards the nascent base pair ^29,32,33^. In both binary1 and 2, the O helix is in a predominantly open position with the N-terminal tip of the O helix rotating towards the active site by only ∼11° (Figure 3A). We will denote this state of the O-helix as quasi-open. Taken together, binary1 and 2 complexes indicate that apPol can engage with the DNA and nucleotide substrates independent of each other.

**Figure 3:**
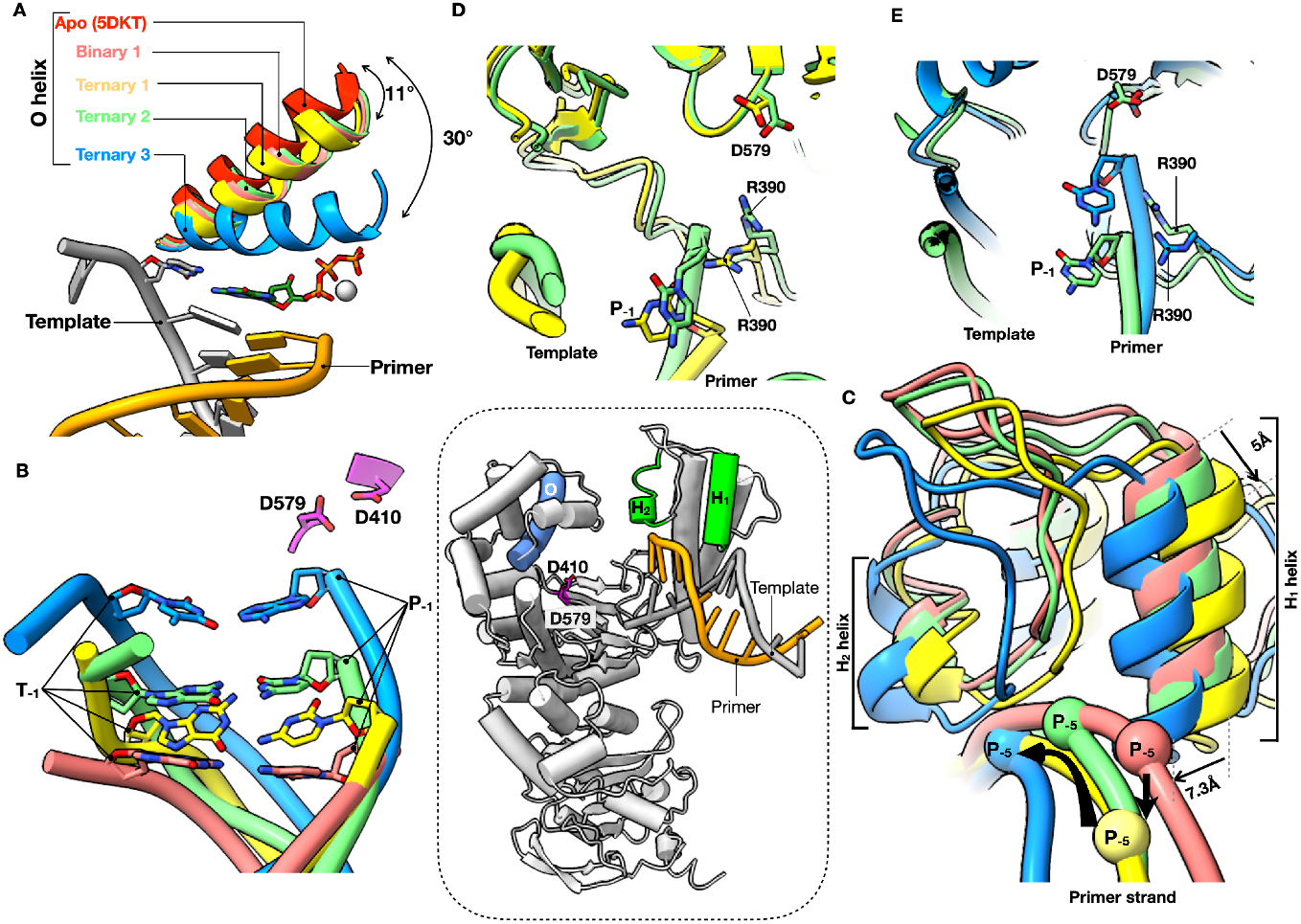
Protein conformation and DNA position changes leading to a catalytically poised apPol complex. **(A)** Position of the O helix (highlighted in blue in the inset) of apPol pre-chemistry complexes with respect to the O helix position of *apo* apPol (PDB id: 5DKT). The primer (orange)/template (grey) DNA, incoming dGTP (green) and Ca^2+^ (white) occupying the position of metal B in ternary3 are shown for reference. The T_0_ base is depicted in stick representation while all other bases of the DNA are shown as slabs. For angle measurement the C_α_ atom of Y471 of *apo* apPol was selected as the vertex. **(B)** Position of the T_-1_-P_-1_ base pair of binary1, ternary1, ternary2 and ternary3 with respect to the catalytic aspartates (D579 and D410; magenta; position of the aspartates highlighted in the inset). The DNA backbones are shown as ribbons with the T_-1_ and P_-1_ bases shown in stick representation. Only the catalytic aspartases of ternary3 are shown for clarity. We note that the peptide backbone of D579 and D410 of all the apPol complexes were aligned to each other. **(C)** Alterations in thumb and DNA positions among the different apPol complexes. For clarity, only H_1_ and H_2_ helices (highlighted in green in the inset) of the thumb along with the connecting loop and the DNA primer strands are shown. The spheres on the primer strands denote the P_-5_ positions and are used to illustrate the DNA motion. Black arrows show the potential thumb (thin arrows) and DNA (thick arrows) motions. **(D, E)** Position of R390 in ternary1 and ternary2 **(D)** and ternary2 and ternary3 **(E)** with respect to the 3’ end of the primer. R390, D579 and the P_-1_ nucleotide are shown in stick representation. Colour coding for all panels: salmon pink: binary1; yellow: ternary1; green: ternary2; blue: ternary3; red: apo. **Inset:** apPol model from Figure 1B with the DNA omitted. The protein in coloured white with the O (blue), H_1_ and H_2_ (green) helices and D410 and 579 (magenta) highlighted.

### apPol-DNA binary complex base pairs with the incoming nucleotide with the fingers open

We solved the structure of apPol-DNA-dGTP ternary complex in three conformational states (ternary1, 2 and 3) (Figure 1C). Ternary1 and 2 were resolved through 3D classification of dataset1 while ternary 3 is from dataset2 (Figure S2). The overall protein structure is similar among these states (average C_α_ RMSD between ternary1 and 2 is 1.5Å and between ternary1 and 3 and ternary2 and 3 are 2.8Å and 2.9Å respectively). In ternary1 and 2 the O helix is quasi-open, while in ternary3 the fingers is in a closed conformation with the O helix rotating towards the active site by 30° (Figure 3A; Movie 1). By comparing the DNA positions and active site configurations of the ternary complexes with those of binary1, we could assign the complexes on a reaction pathway starting with the initial collision dGTP with the apPol-DNA binary complex to the catalytically poised pre-chemistry state.

The ternary1 DNA position is closest to that of binary1 (Figure 3B) and we assign it as the initial collision ternary complex capturing an initial contact of the incoming dGTP with the apPol-DNA pre-chemistry binary complex. Going from binary1 to ternary1 the H_1_ and H_2_ helices of the thumb push down on the DNA such that the DNA pivots around P_-3_, and the T_-1_-P_-1_ base pair swings in deeper into the palm domain getting closer to the catalytic aspartates (D410 and D579) (Figures 3C and B; Movies 2 and 3). In this state the 3’ end of the primer strand is ∼17 Å away from D579 (Figure S4).

We could detect 20 base pairs in the duplex DNA but no density was present for the template overhang except for the T_0_ base (Figure 1C). We could detect the density for the incoming dGTP in between the fingers and palm domains (Figure S3D). The triphosphate moiety of the dGTP is in an extended configuration and is stabilised by Q413 of the fingers domain (Figures 4A and S3D).

**Figure 4:**
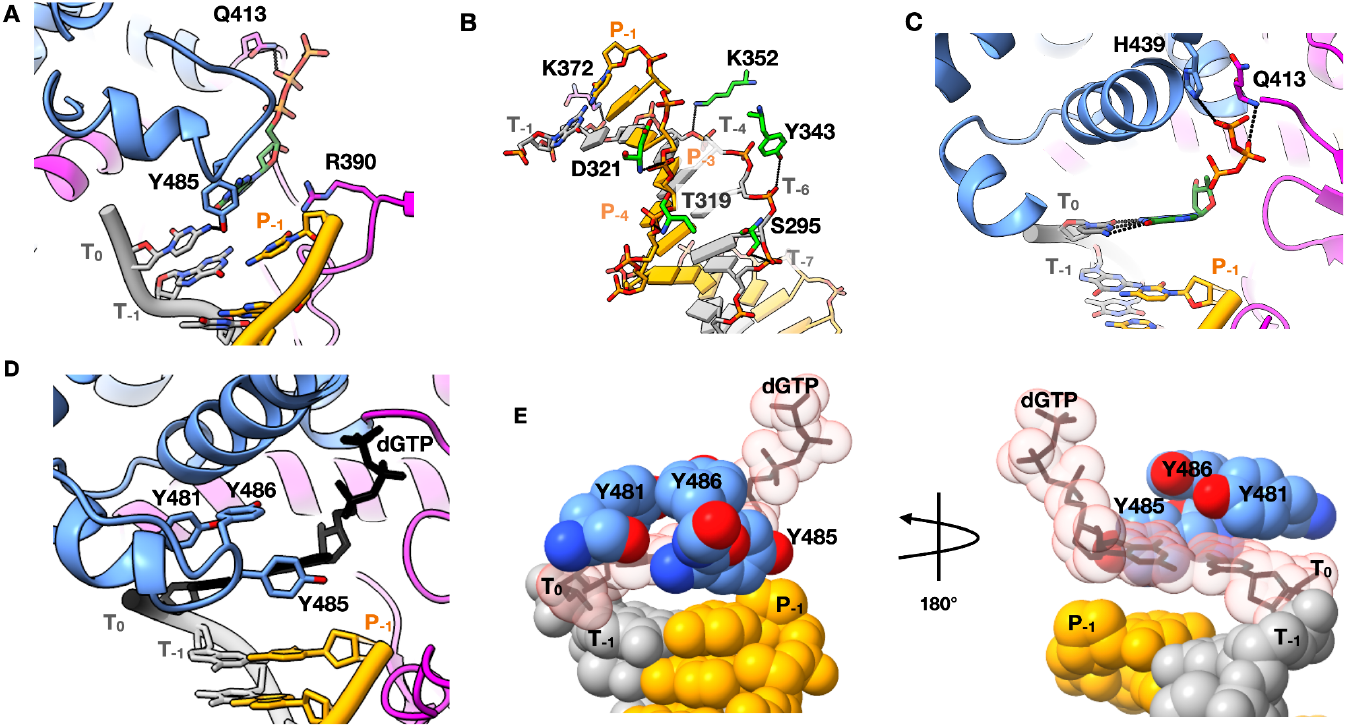
Open ternary complexes of apPol. **(A)** The incoming dGTP in ternary1. Residues interacting with the dGTP and R390 are shown in stick representation. **(B)** Specific interactions of apPol with the DNA duplex in ternary2. The bases are shown as slabs except T_-1_ and P_-1_, which, along with the interacting residues are shown in stick representation. **(C)** The nascent base pair in ternary2. Residues interacting with the dGTP are shown in stick representation. **(D, E)** Pre-insertion checkpoint formed by Y481, Y485 and Y486 of the O1 helix in ternary2. **(D)** Cartoon representation of the pre-insertion checkpoint with the tyrosines shown in stick form. **(E)** The three tyrosines and the DNA duplex are shown as spheres with the incoming dGTP and the T_0_ base shown both in sphere and stick representations. Colour coding for all panels is the same as Figure 1B with the incoming dGTP shown in dark green in panels A and C. The sphere representation of the T_0_ base and the incoming dGTP (panel E) are shown as transparent salmon pink and the corresponding stick representations (panels D and E) are in black.

Compared to the triphosphate, the densities for the ribose and base were weak and we could only tentatively model their orientations. We postulate that in ternary1 the incoming dNTP can sample multiple conformations and once favorable pairing occurs with the T_0_ base the complex moves to ternary2. In addition to the apPol-DNA contacts of binary1, we note a hydrogen bond between Y485 of the O1 helix of the fingers domain and the T_0_ base (Figure 4A). Interestingly, the sidechain of R390 is sandwiched between the 3’ end of the primer and the incoming dGTP and acts as a roadblock for the DNA moving towards the fingers (Figures 4A and S4).

### apPol-specific O1 helix extension acts as a nascent base pair checkpoint

Going from ternary1 to ternary2, R390 adopts an altered conformation, no longer restricting the 3’ end of the primer terminus (Figures 3D, and S4). This altered R390, combined with a rigid body motion of the thumb towards the fingers domain (Figure 3C), guides the DNA towards the fingers and moves the 3’ end of the primer closer to the catalytic aspartates by ∼6Å (Figures 3B, C and S4). In ternary2, the template strand forms multiple hydrogen bonds with the thumb and palm domain residues (Figure 4B). The incoming dGTP pairs with the T_0_ dC and the pairing is consistent with a Watson-Crick base pair (Figure 4C). Y481, Y485 and Y486 of the O1 helix form a snug cavity for the nascent base pair (Figures 4D, E and S3E; Movie 4). Compared to the structurally characterised A-family polymerases, the O1 helix of apPol is longer by ∼1 turn at the C-terminal end (Figure 2C) and Y481, Y485 and Y486, conserved within the apPol clade of A-family polymerases ^24^, are positioned in this additional helical turn. Compared to the catalytically poised closed ternary complexes of DNA polymerases, the triphosphate moiety of the dGTP adopts a different orientation with the oxygens of the gamma phosphate curled away from the catalytic aspartates (Figures 4C and S3E). The dGTP is stabilised by H439 and the backbone of Q413. We could not detect any metal ion density with the incoming dGTP in ternary1 or 2.

### Catalytically poised apPol has the fingers closed around the nascent base pair

To capture ternary3 we used a primer strand with 3’OH but instead of Mg^2+^ we used Ca^2+^ as the divalent metal ion (dataset2) (Figure 1A). It has been shown that while Ca^2+^ allows the formation of the catalytically poised ternary complex, for most polymerases catalysis is prevented in part due to the subtle differences in coordination geometries and the size difference between the calcium and magnesium ions ^8,22,34^. Based on the active site arrangement of ternary3 (Figure 5A), we conclude that this state is the catalytically poised state of apPol.

**Figure 5:**
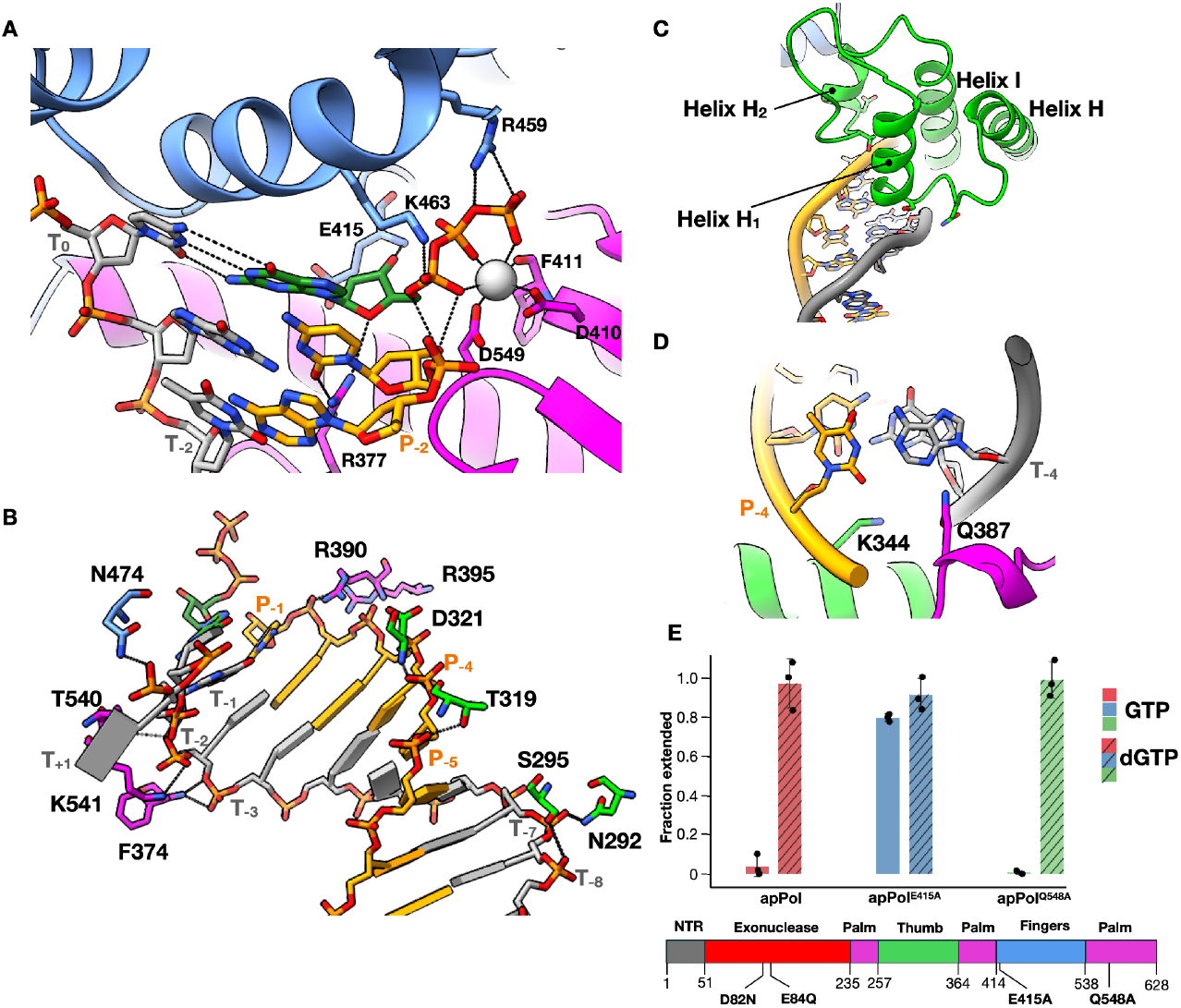
Catalytically poised apPol. **(A)** Polymerisation active site of ternary3. Residues interacting with the incoming dGTP are shown in stick representation. **(B)** Specific interactions of apPol with the DNA duplex in ternary3. The bases are shown as slabs except T_-1_ and P_-1_. These bases, incoming dGTP and the interacting residues are shown in stick representation. **(C)** Thumb elements of ternary3 interacting with the minor groove backbone of the primer/template DNA. The residues involved in specific contacts are shown in stick representation. **(D)** Top-down view of the primer/template DNA duplex in highlighting Q387 and K344 sensing the minor groove geometry of the T_-4_-P_-4_ base pair. Colour coding in panels A, B, C and D is the same as Figure 1B with the incoming dGTP shown in dark green and Ca^2+^ is represented as white sphere. **(E)** Single nucleotide primer extension assay to validate the steric gate residue. 1μM apPol (either apPol (Figure 1B) or apPol with an additional point mutation (E415A or Q548A)) was incubated with 50nM FAM-P/T. 125μM of dGTP (hashed bars) or GTP (solid bars) was added to initiate the reaction. The reactions were quenched after 2 minutes with 250 mM EDTA and the fraction of starting primer extended by one nucleotide has been plotted as bars. The experiments were performed in triplicates and the individual replicates are shown as black circles with the average value shown as bars. Error bars show the standard deviation.

Going from ternary2 to ternary3, the thumb undergoes a rigid body motion towards the fingers, leading to a stronger interaction with the DNA and corkscrew movement of the DNA into the apPol active site (Figures 3C, B and S4; Movies 2, 3 and 5). The DNA duplex in ternary3 is stabilised through extensive contacts with apPol (Figure 5B) with a total buried surface area of 1798Å^2^. In this state the thumb tracks both the backbone geometry and the base pair orientation of the DNA minor groove (Figures 5C and D; Movie 2) and this might act as a checkpoint for potential misincorporations. In ternary3 the 3’ primer terminal base is stabilised by a minor groove hydrogen bond with R377 (Figure 5A), placing the P_-1_ C3’ 4.2Å away from alpha phosphate (P_?_) of the incoming dGTP. Notably, the position of the 3’ end of the primer strand of ternary3 is occupied by R390 in ternary2 (Figure 3E). Thus, as the DNA moves into the active site, R390 adopts an altered conformation to accommodate the primer strand (Figures 3E and S4).

In ternary3 the O helix of the fingers domain clamp down on the nascent base pair, restricting the active site of apPol in preparation for chemistry (Figures 3A and 5A) . The closure of the O helix is accompanied with a concomitant movement of the O1 helix away from the catalytic aspartates (Movies 1 and 4). The O helix closure pushes the dGTP down into the active site with the triphosphate in a configuration conducive for catalysis (Figure 5A). The triphosphate is stabilised through interactions with R459 and K463 of the fingers. We could detect the density for metal B, coordinated by the two catalytic aspartates (D579 and 410), F411 backbone and the triphosphate of the incoming dGTP (Figure 5A). Interestingly, we could not detect any density for metal A, indicating that metal A might have weaker binding affinity compared to metal B. Instead, we find the 3’OH of the primer strand to form hydrogen bonds with the incoming dGTP. We interpret the ribose pucker of the incoming nucleotide as C3’-endo (Figure S5A), similar to the sugar pucker observed in other catalytically poised DNA polymerases. From the ternary3 structure we identify E415 as the steric gate residue ^35^. The C2’ of the dGTP is positioned 3.7Å above E415 (Figure 5D). We anticipate that this residue will discriminate against an incoming NTP through steric clash with the 2’OH. In fact, this glutamic acid is conserved in A-family polymerases and acts as the steric gate ^7,35–37^. We performed single nucleotide primer extension assays with both wild type (exonuclease inactivated) and a mutant version of apPol where E415 was mutated to alanine (apPol^E415A^) with either dGTP or GTP as the incoming nucleotide (Figures 5E, S5B and C). We found that, unlike apPol, apPol^E415A^ can incorporate both GTP and dGTP opposite dC (Figure 5B), although GTP incorporation was slower than dGTP(Figures S5B and C). This indicates that E415 indeed plays a key role in discriminating GTP over dGTP as the incoming nucleotide. Recently it has been proposed that Q548 might act as the steric gate residue for apPol ^27^. However, our primer extension assay shows that Q548 does not discriminate against GTP (Figures 5E and S5C).

Ternary3 is the only state where we could detect template overhang density for the T_+1_ position (Figure 5B). N474 forms a hydrogen bond with the phosphate of T_+1_ thus making the base ordered. The T_+1_ base is in a flipped-out configuration, excluding it from the active site. This flipped-out configuration is essential to avoid a steric clash with Y481.

## DISCUSSIONS

The series of cryoEM structures reported here (Figure 6) captures how apPol transitions through the initial stages of interactions with its DNA and nucleotide substrates, ultimately forming a catalytically poised pre-chemistry ternary complex while maintaining Watson Crick geometry of the nascent base pair. We discuss our results within the context of A-family polymerases and propose a mechanism by which apPol might accommodate DNA lesions within its active site while biassing the nascent base pair towards Watson Crick geometry, a prerequisite for performing both replicative and lesion bypass synthesis within the same DNA polymerase.

**Figure 6:**
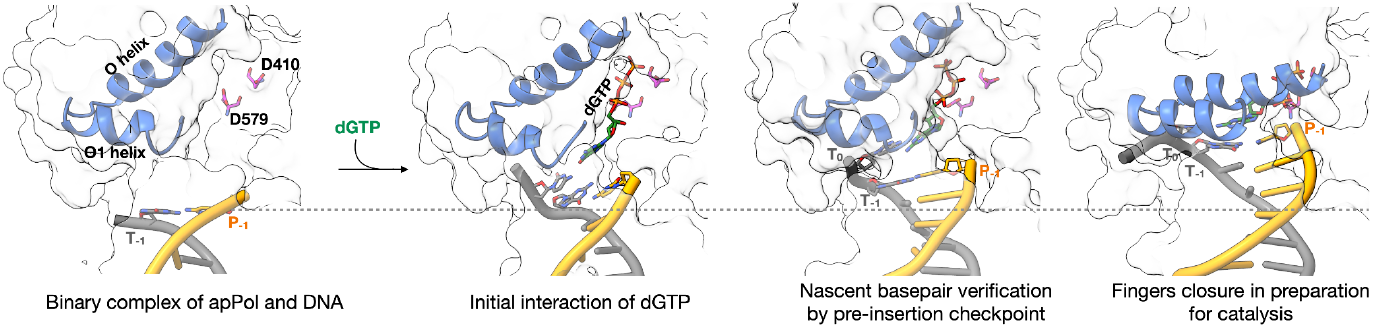
apPol backtracks during the pre-chemistry steps. Zoomed-in views of 3’ primer terminus in binary1, ternary1, ternary2 and ternary3 complexes (left to right) with the DNA primer (orange)/template (grey), O and O1 helices (blue) shown in ribbon representations. The dGTP (green), catalytic aspartates (magenta) and the P_-1_, T_-1_ and T_0_ bases are shown in stick representations. The rest of the apPol is shown as a translucent grey surface. The grey dashed line marks the position of P_-1_ base in binary1. All structures were aligned on the palm domain.

### apPol uses two fidelity checkpoints to accommodate TLS and replicative synthesis

DNA polymerase I has been a model A-family replicative polymerase and, based on a series of structures of *Bacillus stearothermophilus* polymerase I (Bst Pol I) performing nucleotide addition *in crystallo*, a conformational coupling mechanism was proposed for high fidelity replication ^38–40^. In the pre-chemistry binary complex the O helix remains open with the insertion site (position of T_0_ base during catalysis) occupied by a tyrosine residue at the C-terminus of the O helix (Y714 in BSt Pol I) (Figure 2C). This tyrosine is conserved in all A-family polymerases and occupies the insertion site in most apo or binary A-family polymerase structures ^33,41–43^, potentially preventing premature base pair formation with the fingers in the open configuration. In the fingers open state of Bst Pol I, the T_0_ base remains lodged within a pocket formed by the O and O1 helix (pre-insertion site) (Figure 2C). As the polymerase moves from binary to ternary state, the O helix closes, leading to a conformational change in the loop connecting the O and O1 helices, which (a) blocks the pre-insertion site and (b) alters the orientation of Y714 and freeing up the insertion site. Thus the T_0_ base is forced out of the pre-insertion site and into the newly available insertion site. Moreover, the altered conformation of the loop connecting the O and O1 helices forces the T_+1_ base to adopt a flipped-out orientation to avoid any steric clashes. These coupled conformational changes ensure that (a) no premature bond formation occurs with the fingers open, thus biasing the nascent base pair towards Watson-Crick geometry (since in the closed state the active site can accommodate predominantly Watson-Crick pairing) and (b) only a single templating base gets access to the insertion site during a single round of the catalysis, thereby reducing the chance of a frameshift error.

ApPol does not undergo the coupled conformational changes noted for Pol I. In the binary1 complex the pre-insertion site is blocked by M473 and Y481, excluding the T_0_ base from this position (Figures 2C and S3B). Moreover, the tyrosine at the C terminus of apPol’s O helix (Y471) does not occupy the insertion site (Figures 2C and S3B) and consequently apPol can form the nascent base pair with the fingers in the quasi-open state as seen in ternary1 and 2 (Figures 4A and C; Movie 1). This would allow apPol to accommodate a nascent base pair that deviates from the Watson-Crick geometry, a feature that might aid in apPol’s TLS activity.

This raises the question how does apPol ensure fidelity during nucleotide incorporation? We hypothesize that apPol uses two fidelity checkpoints. The first checkpoint is captured by ternary2 where the nascent base pair fits within the snug pocket formed by Y481, Y485 and Y486 at the tip of the O1 helix (Figures 4D, E and S3E; Movie 4). This pocket is in fact a modified pre-insertion site. By modifying this site, apPol has converted the pre-insertion site into a pre-insertion checkpoint to assess the nascent base pair geometry. Notably, the residues involved in forming the snug pocket (Y481, Y485 and Y486) are conserved in the apPol clade of the A-family but absent in other A-family polymerases studied to date ^26^. Consistent with the critical role of these three residues, it has been reported that a mutant version of apPol where these three tyrosine are mutated to alanines shows severe defects in primer extension activity ^26^. The second checkpoint is captured by ternary3 where the nascent base pair is poised for catalysis and the fingers is in a closed conformation (Figure 5A). In this state the active site will favour the Watson-Crick base pair geometry. These two fidelity checkpoints might allow apPol to perform TLS without sacrificing the accuracy of replicative synthesis.

### apPol thumb is a master manipulator of DNA position

In all A family polymerases studied to date, the DNA undergoes minimal movement between the binary and ternary complexes ^33,38,43^. However, apPol is an exception to the rule with the enzyme backtracking with respect to the DNA ^44^ during transition from pre-chemistry binary to the catalytically poised pre-chemistry ternary state (Figure 6; Movie 3). Going from binary1 to ternary1 the DNA moves towards the active site facilitated by a downward motion of the thumb (Figure 3C; Movie 2), pushing the 3’ end of the primer towards the active site. Further, as apPol transitions from ternary1 to ternary2 and finally to ternary3 the DNA undergoes additional corkscrew motion positioning the 3’ end of the primer into the active site (Figures 3B and S4; Movie 3).

This large scale DNA movement is orchestrated in part by a rigid body motion of the thumb towards the fingers domain. In addition, the DNA slips with respect to the thumb to achieve the full extent of the movement. For instance, T319 of the thumb domain contacts the phosphate backbone of P_-3_ in binary1 and ternary1, P_-4_ in ternary2 and P_-5_ in ternary3 (Figures 2B, 4B and 5B; Movie 6). As apPol progresses from binary1 to ternary3 the template strand forms contacts with residues from the palm and fingers and with the incoming nucleotide (Movie 5, Figures 5A and B); alternating breakage and re-formation of the contacts with the thumb and residues in the vicinity of the active site leads to the DNA slippage.

Unlike other A-family polymerases, the thumb of apPol makes few contacts with the DNA, especially in the binary state, and this might explain the relatively weak affinity of DNA for apPol ^5,27^. For instance, Bst Pol I and T7 DNA polymerase binary complexes have polymerase-DNA interfaces of 1782Å^2^ (PDB id: 1L3T) and 2009Å^2^ (PDB id: 2AJQ) respectively while the corresponding interface for apPol is only 607Å^2^. For the apPol-specific pre-insertion checkpoint to function, the DNA and the incoming nucleotide needs to undergo large scale motions as apPol transitions from the pre-chemistry binary complex to the catalytically poised complex. The weaker contact between the apPol thumb and the DNA might have evolved to facilitate the large DNA motion enabling this apPol-specific checkpoint.

### Sugar pucker of the incoming dGTP

Recently, a pre-chemistry ternary complex structure of apPol with the fingers in closed configuration has been published (referred to as ternary4 from here onwards) ^27^. The overall conformation of apPol is similar between ternary3 and ternary4, with the main difference being the orientation of the O helix. While the O helix of ternary4 is almost closed, the closure is not complete (Figure S5D). On the other hand there are some critical differences in the orientation of the incoming dGTP. In ternary4, unlike other DNA polymerase structures that we are aware of, including ternary3, the ribose sugar of the incoming dGTP has been modelled with an O4’-endo sugar pucker (Figure S5A). The altered pucker in ternary4 compared to ternary3 alters the orientation of the incoming deoxyribose with respect to the 3’ end of the primer (Figure S5E) and results in several steric clashes including a clash between C2’ of P_-1_ and C1’ of the incoming dGTP. It is puzzling why a matched incoming nucleotide (dGTP opposite dC in ternary4) would adopt an unusual sugar pucker. The published ternary4 structure was at an overall resolution of 3.2Å. At this resolution it is challenging to unambiguously determine the sugar pucker and higher resolution structures of ternary4 would be essential before any further analysis is possible.

### Why do the fingers close?

In most catalytically poised DNA polymerase ternary complexes, including apPol, the fingers domain stays in a closed conformation ^6,8,29^. On the other hand, in the apo or binary states, the fingers are typically open. Closed fingers restrict the polymerisation active site and sequesters the 3’OH of the primer and incoming nucleotide from the bulk solvent, facilitating the phosphodiester bond formation. Nucleotide binding has typically been assigned as the trigger for fingers closure ^45^.

However, it has been shown that the Klenow fragment of *E. coli* Pol I and the herpes simplex virus-1 DNA polymerase can sample both the open and closed conformations in the absence of the incoming nucleotide ^28,42,46^. This indicates that dNTP binding is not essential for fingers closing. It is possible that the closed state is stabilised in the presence of the correct incoming nucleotide and/or the two divalent metal ions associated with the nucleotide ^28,47,48^. However, recent structures of the human leading strand DNA polymerase, polε, shows that the pre-chemistry ternary complex with the correct nascent base pairing can exist with the fingers in open, closed and intermediate states ^49^, indicating that even in the presence of the correct nucleotide, the fingers may stay open. These new observations raise the question, what triggers fingers closing. A combination of kinetic and time-resolved structural measurements investigating the short lived pre-chemistry states of a DNA polymerase would be essential to answer this question, which is central to our understanding of the catalytic mechanisms of DNA polymerases.

### What happens after bond formation?

The series of structures presented here elucidate how apPol interacts with DNA and the incoming nucleotide to catalyse phosphodiester bond formation; however we do not have direct structural insight into the post-chemistry events. While a post-chemistry binary complex (Figure S1A, step 4) structure of apPol (generated through addition of a dideoxyATP to the 3’ end of the primer strand) has been reported ^27^, the modest resolution of 5.2Å precludes drawing any atomistic inference. In fact, at this resolution, it is difficult to ascertain whether the structure is of a post-chemistry binary state or whether apPol has translocated (Figure S1A, step 5) to a pre-chemistry binary form.

Nonetheless, based on the DNA position in binary1 and the published post-chemistry binary structure we predict that after chemistry apPol over-translocates leading to the apPol-DNA pre-chemistry binary complex (binary1). Following this apPol backtracks to correct for the over-translocation and positions the templating DNA base in the active site.

Among the well-studied replication systems, the T7 bacteriophage replisome is perhaps the closest to the apicoplast replisome ^50^. *In vitro* studies indicate that in T7 replisome the leading strand gp5 T7 DNA polymerase follows the gp4 helicase-primase closely with only a single nucleotide of the DNA template separating the two proteins ^51,52^. However, the over-translocation mechanism for apPol would require a larger slack in the DNA between apPol and the apicoplast helicase or any other partner protein positioned downstream of the polymerase. Alternatively, the over-translocation might be used by apPol only under special situations like lesion bypass and during bulk DNA synthesis a close contact with a partner protein like the helicase might prevent apPol from over-translocating. Understanding the molecular mechanism of apPol’s post-chemistry substrate reorientation and its implications for the apicoplast replisome organisation would be vital for developing a clear picture of how the apicoplast genome is copied.

## METHODS

### DNA substrates

All DNA oligonucleotides were purchased from Integrated DNA Technology (USA). For dataset1, the primer strand was synthesized with a dideoxy-C at the 3’ terminus. Primer/template DNA substrates were generated by annealing the primer and template DNA strands (sequences shown in figure 1A). All annealing reactions were performed using a 1:1.1 ratio of primer and template DNA in annealing buffer (10 mM Tris-Cl (pH 7.5) and 50 mM NaCl). The sample was heated to 95°C for 5 minutes, followed by gradual cooling to 25°C.

### Overexpression and purification of apPol

For both the datasets, we have used a construct of apPol with two point mutations (D82N and E84Q) at the active site of the proofreading exonuclease domain (Figure 1B). This construct has been referred to as apPol throughout the manuscript. For primer extension assays to determine the steric gate residue, an apPol construct with an additional point mutation (E415A or Q548A) was used. All the constructs had an N-terminal hexa-histidine tag followed by a tobacco etch virus (TEV) protease cleavage site. The constructs were codon optimized for expression in *Escherichia coli*, and were synthesized and cloned into the pETDuet1 vector by GenScript Corp. (USA).

All apPol constructs were overexpressed and purified following a common protocol ^5^. The plasmid was transformed into Rosetta 2(DE3) *E. coli* cells (Merck, USA). The cells were grown in autoinduction terrific broth (Formedium) at 37 °C for ∼5 hours, followed by a further 20 hours of growth at 20°C. The subsequent steps were performed at 4°C. Cell pellets were resuspended in lysis buffer (50 mM Tris–HCl (pH 7.5), 800 mM NaCl, 25 mM imidazole, and 10% glycerol). To prevent proteolytic degradation of apPol and improve lysis efficiency, ethylenediaminetetraacetic acid (EDTA)-free protease inhibitor tablet (Roche, USA) and lysozyme were added to the resuspended cells. Cells were lysed by sonication and clarified by centrifugation. The clarified lysate was loaded onto a 5 ml HiTrap Chelating HP column (Cytiva, USA) charged with Ni^2+^ and pre-equilibrated with lysis buffer. The unbound protein was washed with 10 column volumes (CVs) of lysis buffer, followed by an additional 10 CV wash with low-salt wash buffer (50 mM Tris–HCl (pH 7.5), 200 mM NaCl, 25 mM imidazole, and 10% glycerol). ApPol was eluted over a 10 CVs linear gradient of imidazole from 25 mM to 1 M. The elution fractions were analyzed using SDS-PAGE, and fractions containing apPol were pooled and diluted with no-salt buffer (50 mM Tris–HCl (pH 7.5), 25 mM imidazole, and 10% glycerol) to decrease the NaCl concentration to 100 mM. The diluted sample was loaded on a 5 ml HiTrap SP column (Cytiva, USA) pre-equilibrated with Buffer A (50 mM Tris–HCl (pH 7.5), 100 mM NaCl, 5 mM EDTA, 5 mM β-mercaptoethanol, and 10% glycerol). Unbound protein was washed with 10 CVs of Buffer A, and apPol was eluted with a linear gradient of 0.1 to 1 M NaCl (10 CVs). For further purification, the eluent of the cation exchange column was concentrated and loaded on a Superdex 200 Increase 10/300 size-exclusion chromatography column (Cytiva, USA) pre-equilibrated with storage buffer (50 mM Tris-Cl (pH 7.5), 200 mM NaCl, 2 mM DTT, and 20% glycerol). Fractions containing pure apPol were pooled, concentrated, flash-frozen in liquid nitrogen, and stored at −80°C. Protein concentration was determined based on the theoretical extinction coefficient of 64180 M^−1^ cm^−1^.

The hexa-histidine tag was not removed prior to using the protein.

### Primer extension assays

All primer extension assays were performed at 37°C in reaction buffer containing 20 mM Tris-Cl (pH 7.5), 10 mM MgCl_2_, 30 mM NaCl, 1 mM DTT, 0.1 mg/mL BSA, and 5% glycerol. A multiple nucleotide primer extension assay was performed to verify apPol’s activity in presence of 8mM CHAPSO. A final concentration of 480nM apPol was incubated with 200nM primer/template DNA (Figure 1A) with the primer strand having a FAM label at the 5’ end (FAM-P/T). The reaction was initiated by adding 250 μM of each dNTP. The reaction was quenched after 10 or 20 seconds by adding 250 mM EDTA.

Single nucleotide primer extension assays were performed to ascertain the identity of the steric gate residue of apPol. A final concentration of 1μM apPol (or apPol^E415A^ or apPol^Q548A^) was incubated with 50nM FAM-P/T and 125μM of dGTP or GTP was added to initiate the reaction. The reactions were incubated for varying time intervals ranging from 0 to 120 seconds and then quenched with EDTA.

The products (primer strand extended by multiple or a single nucleotide) were analysed on 15% polyacrylamide (19:1) urea gels run at 40-45°C. The gels were imaged on a Typhoon FLA7000 LASER-based scanner (Cytiva, USA) using an excitation wavelength of 488 nm (blue LASER) and an emission cutoff at 525 nm to detect the fluorescence signal from the 5’ FAM label of the primer strand. For single nucleotide primer extension assays, gel bands were quantitated using ImageQuant software (Cytiva, USA), and the fraction of DNA extended was calculated using the following equation.

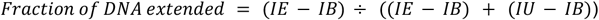

where *IE* is the intensity of the extended primer band, *IB* is the intensity of the gel background, and *IU* is the intensity of the unextended primer band. The data were plotted using RStudio (Posit). A detailed list of R packages used for graph plotting is provided in Table S1.

### CryoEM sample preparation

For dataset1, a ternary complex of apPol, DNA and dGTP was prepared by incubating 20μM apPol with 30μM primer (3’dideoxy-terminated)/template DNA (Figure 1A) and 6mM dGTP at 37°C for 10 minutes in complex-formation buffer (20 mM Tris-Cl (pH 7.5), 10 mM MgCl_2_, 65 mM NaCl and 1 mM DTT). The ternary complex for dataset2 was formed similar to dataset1 with the following three alterations. First, the primer strand was not 3’ dideoxy terminated; second, the complex-formation buffer had 10mM CaCl_2_ instead of 10 mM MgCl_2_; and third, 4mM of dGTP was added instead of 6mM.

8mM CHAPSO was added to the samples immediately before grid preparation. CryoEM grids were prepared using a Leica EM GP automatic plunge freezer (Leica Microsystems). 4μL of the sample was applied to glow discharged Quantifoil R1.2/1.3 400 mesh holey carbon grids (SPT Labtech).

Excess liquid was blotted for 2 seconds and the grids were immediately plunge-frozen in liquid ethane. The vitrified samples were stored in liquid nitrogen until imaged.

### CryoEM imaging and 3D reconstruction

CryoEM grids were screened on a FEI Tecnai Arctica electron microscope (FEI, USA) and the final high resolution datasets were collected on a Titan Krios (ThermoFisher Scientific) operated at 300 kV. Dose fractionated movies were recorded on a K3 direct electron detector (Gatan Inc.) operated in super-resolution mode. Data collection was automated using EPU (ThermoFisher Scientific) and the images were collected at a nominal magnification of 81,000X such that the object level pixel size was 1.06 Å/pixel (super-resolution pixel size: 0.53 Å/pixel). The images were recorded as 1 second movies divided into 70 frames. The total dose and fluence were 70 electron/Å^2^ and 1 electron/Å^2^/frame respectively for dataset1 and 60 electron/Å^2^ and 1.2 electron/Å^2^/frame respectively for dataset2.

All image processing jobs and three-dimensional (3D) reconstructions (Figure S2) were performed using CryoSPARC (versions 4.5.1) ^53^ (Structura Biotechnology Inc.). The individual super-resolution movie frames were binned by 2 and the frames were aligned using alignparts_lmbfgs ^54^ as implemented within Cryosparc. The contrast transfer function (CTF) of the micrographs were estimated using the patch CTF routine of Cryosparc.

For dataset1, after removing unsuitable micrographs (micrographs with defocus higher than -3.8 μm, 0.5 CTF fit resolution worse than 6 Å, and relative ice thickness higher than 1.1 were rejected), 14,519 micrographs were retained out of 16,197 for further processing. Particles were picked using Topaz ^55^. The training set contained ∼1400 manually picked particles from micrographs with varying defocus. At this stage, 1,922,870 particles were selected. These particles were subjected to multiple rounds of 2D classification and 898,336 particles contributing to optimal 2D class averages were retained for further processing. These particles were used for reference-free 3D reconstruction using Ab-initio reconstruction ^53^. One of the classes (354,534 particles) showed clear features consistent with apPol and was taken forward for further processing. These particles were refined using the non-uniform refinement routine ^56^ resulting in a consensus map with a nominal resolution of 3.2Å. Multiple rounds of 3D classification (without alignment) were done including masked classification ^57^ focussing on the DNA and nucleotide substrates to resolve the heterogeneity in the dataset. This resulted in four different classes defined as binary1, binary2, ternary1 and ternary2. Residual beam-induced motion and defocus estimation errors of the particles contributing to these classes were rectified using the reference based motion correction and per particle CTF estimation routines of Cryosparc. This was followed by 3D refinement (using the non-uniform refinement routine of Cryosparc) of each of the 3D classes resulting in the final maps.

For dataset2, after removing unsuitable micrographs (micrographs with average defocus greater than -3.8μm, 0.5 CTF fit resolution worse than 5 Å, full frame motion greater than 27.4 pixels and relative ice thickness greater than 1.0 were rejected). 11,231 micrographs were retained out of 11,944 for further processing. Particles were picked using Topaz. The training set contained ∼1400 manually picked particles from micrographs with varying defocus. At this stage, 1,993,221 particles were selected. These particles were subjected to multiple rounds of 2D classification and 238,125 particles contributing to optimal 2D class averages were retained for further processing. These particles were used for reference-free 3D reconstruction using Ab-initio reconstruction and one class resembling apPol (102,768 particles) was taken forward for further processing. Multiple rounds of heterogeneous refinements were performed to further eliminate particles that do not contribute to high resolution reconstruction. and 92,787 particles were taken forward. The residual beam induced motion was rectified and the defocus of these particles were re-estimated using the reference based motion correction and per particle CTF estimation routines of Cryosparc respectively. This was followed by 3D refinement (using non-uniform refinement) leading to a 3.5Å map of the ternary3 complex.

The resolutions of all cryoEM maps are reported based on the 0.143 cutoff of the gold-standard Fourier shell correlation (FSC) curves ^58,59^. For better map interpretability, the refined maps were sharpened either by applying a negative B factor ^60^, estimated during non-uniform refinement, or using DeepEMhancer ^61^. For DeepEMhancer based sharpening, the tightTagret model was used.

### Atomic model building, refinement and structure analysis

The published apo apPol structure ^26^ (PDB ID:5DKT) was used as the starting model for apPol. The starting model for the primer/template DNA was generated from the structure of *B. stearothermophilus* DNA polymerase1 (BSt Pol I) (PDB id: 1LV5) ^38^ fitted into the sharpened cryoEM maps using the ‘Fit in Map’ tool in ChimeraX ^62^ . The fitted models were examined in Coot ^63^ and fits of side chain rotamers and main chain were optimized either by real space refinement within Coot or in Isolde ^64^ and subjected to real space refinement within Phenix ^65^ . Ligand restraint for dGTP was generated using eLBOW ligand restraint generation ^66^ in Phenix and was used in all dGTP refinement. The refined structures were analysed in Coot and rebuilt where necessary to optimize model geometry (assessed by MoProbity ^67^ ) and the fit of the model within the cryoEM map (Table S2). To estimate over-fitting, FSC curves were calculated between the cryoEM map and the atomic model (map-to-model FSC ^60^ ) in Phenix.

All maps and models were visualised in UCSF ChimeraX and unless otherwise stated, the map density depicted in the figures correspond to the maps sharpened using DeepEMhancer. Map segmentation was performed using Seggar ^68^. For comparison between different apPol complexes, residues 405-411 and 572-585 of apPol were used for alignment. The movies depicting morphs between the different states of apPol were prepared in Pymol (Schrodinger).

## ACKNOWLEDGEMENT

This work was supported by a DBT/Wellcome Trust India Alliance Fellowship [IA/I/20/1/504905] and a Royal Society Research Grant [R/179278] awarded to I.L. and NIH grant R01 GM-080573 to J.D.P. A.K. and T.E. are supported by BBSRC DTP studentships awarded to T.D.C and I.L. [BB/T007222/1]. T.D.C was supported by BBSRC [BB/W00061X/1]. CryoEM grids were screened at the University of Sheffield cryoEM facility and we thank Dr. Svetomir Tzokov for support with this screening. High resolution cryoEM data was collected at the Electron Bio-Imaging Centre (eBIC) [Block Allocation Group number: BI34172] and we thank Dr. Peter Harrison for support with data collection. We thank Drs. Purba Mukherjee (University of York, UK) and James Reid (University of Sheffield, UK), and Professors Sarah Harris (University of Sheffield, UK) and Per Bullough (University of Sheffield, UK) for helpful comments and suggestions at various stages of the work.

We thank all the members of Lahiri and Pata labs for their helpful comments and suggestions about the work.

## CONTRIBUTIONS

Conceptualization, A.K., T.D.C and I.L.; Investigation, A.K., T.E. and I.L.; Formal Analysis, A.K., T.E., J.D.P and I.L.; Resources, I.L.; Writing – Original Draft, A.K and I.L.; Writing – Review & Editing, A.K., T.E., T.D.C, J.D.P. and I.L.; Visualization, A.K., T.E., J.D.P. and I.L.; Supervision, T.D.C. and I.L.; Project Administration, I.L.; Funding Acquisition, T.D.C., J.D.P. and I.L.

## DECLARATION OF INTERESTS

The authors declare no competing interests.

## Notes

### Competing Interest Statement

The authors have declared no competing interest.

