## Supplemental figures and tables for "Structural basis of multitasking by the apicoplast DNA polymerase from *Plasmodium falciparum*"

### Supplementary figures and tables

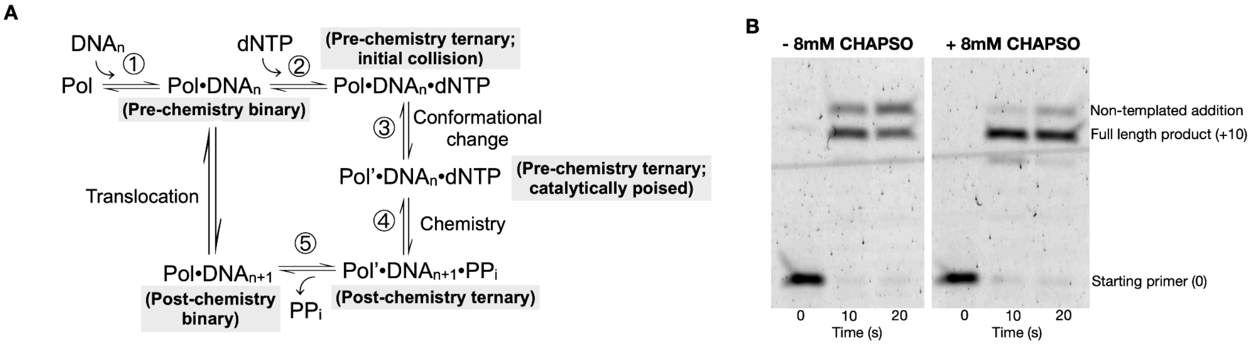


**Figure S1: (A)** Catalytic cycle for nucleotide (dNTP) incorporation by a DNA polymerase (Pol) during processive synthesis. The names of the complexes along the reaction pathway are highlighted in grey. $DNA_n$: DNA substrate with a primer strand n bases long, $DNA_{n+1}$: DNA substrate with a primer strand n+1 bases long, Pol': Catalytically poised DNA polymerase. $PP_i$: inorganic pyrophosphate. **(B)** Multiple nucleotide incorporation by apPol in the absence (left) and presence (right) of 8mM CHAPSO.

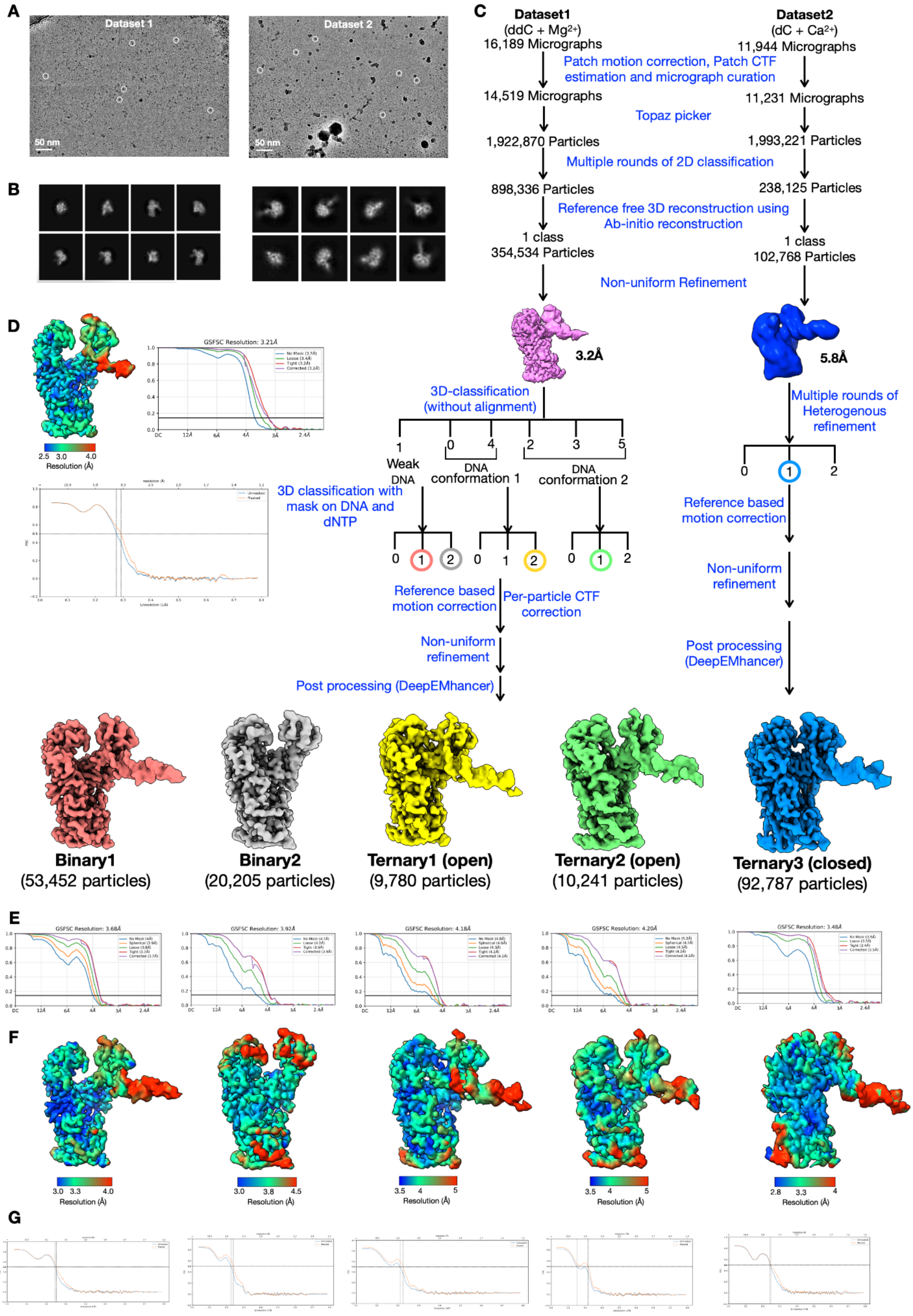

A
Dataset 1
Dataset 2
50 nm
B
C
Dataset1
(ddC + Mg2+)
16,189 Micrographs
Dataset2
(dC + Ca2+)
11,944 Micrographs
Patch motion correction, Patch CTF estimation and micrograph curation
14,519 Micrographs
11,231 Micrographs
Topaz picker
1,922,870 Particles
1,993,221 Particles
Multiple rounds of 2D classification
898,336 Particles
238,125 Particles
Reference free 3D reconstruction using Ab-initio reconstruction
1 class
354,534 Particles
1 class
102,768 Particles
Non-uniform Refinement
3.2Å
5.8Å
3D-classification (without alignment)
Multiple rounds of Heterogenous refinement
Weak DNA
DNA conformation 1
DNA conformation 2
3D classification with mask on DNA and dNTP
Reference based motion correction
Per-particle CTF correction
Non-uniform refinement
Post processing (DeepEMhancer)
D
GSFSC Resolution: 3.21Å
Resolution (Å)
Binary1
(53,452 particles)
Binary2
(20,205 particles)
Ternary1 (open)
(9,780 particles)
Ternary2 (open)
(10,241 particles)
Ternary3 (closed)
(92,787 particles)
E
GSFSC Resolution: 3.68Å
GSFSC Resolution: 3.92Å
GSFSC Resolution: 4.18Å
GSFSC Resolution: 4.20Å
GSFSC Resolution: 3.48Å
F
Resolution (Å)
G

**Figure S2: CryoEM image processing and 3D reconstruction from datasets1 and 2. (A)** Representative micrographs from dataset1 (left) and dataset2 (right) with representative particles highlighted with white circles. **(B)** Representative 2D class averages dataset1 (left) and dataset2 (right). **(C)** Schematic representation of the image processing pipeline for datasets1 and 2. The post processed maps are coloured as follows: binary1 (salmon pink), binary 2 (grey), ternary1 (yellow), ternary2 (green) and ternary3 (blue). Coloured circles: classes that were taken forward along the pipeline. Colour coding of the circles is the same as that of the maps. **(D)** The 3.2Å consensus map of dataset1 (pink in panel C) coloured according to local resolution (top left). Gold standard FSC curve of the consensus map (top right) and FSC curve between the consensus map and the corresponding atomic model (bottom). **(E, F, G)** Gold standard FSC curves **(E)**, maps coloured according to local resolution **(F)** and FSC curve between the cryoEM maps and the corresponding atomic models **(G)** of binary1, binary2, ternary1, ternary2 and ternary3 (left to right).

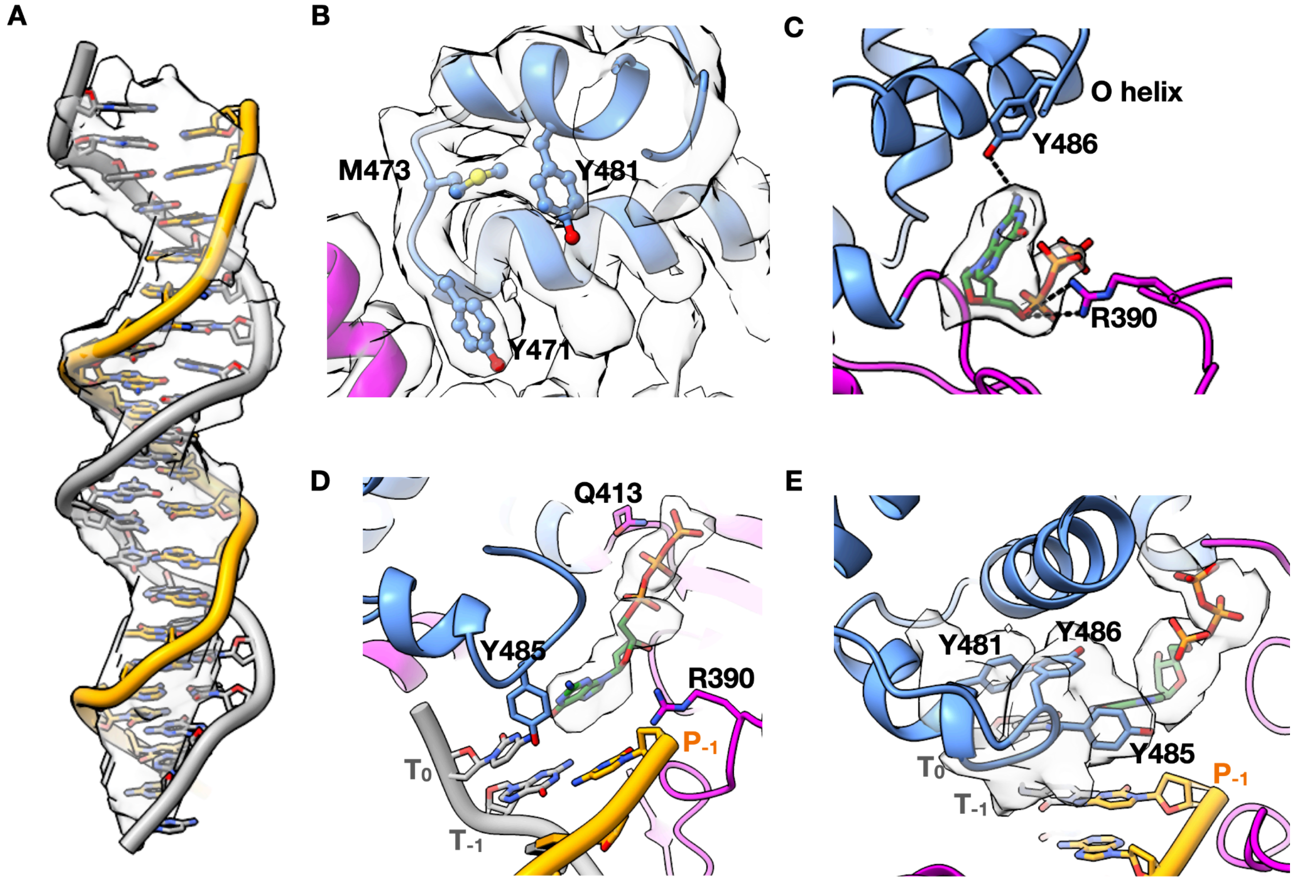


**Figure S3: Fit of model in cryoEM maps. (A)** Primer/template DNA of binary1 fitted into the cryoEM density for the DNA segmented out from the unsharpened map. **(B)** Atomic model of binary1 in the orientation shown in Figure 2C with the sharpened cryoEM map (transparent grey) superimposed on it. **(C)** Interaction of dGTP with apPol in binary2. The cryoEM density for the nucleotide is shown in transparent grey. **(D)** View of ternary1 from Figure 4A with the cryoEM density of the incoming dGTP shown in transparent grey. **(E)** CryoEM map density (transparent grey) around Y481, Y485, Y486 and the incoming dGTP in ternary2. Colour coding for all panels is the same as Figure1B with the incoming dGTP shown in dark green.

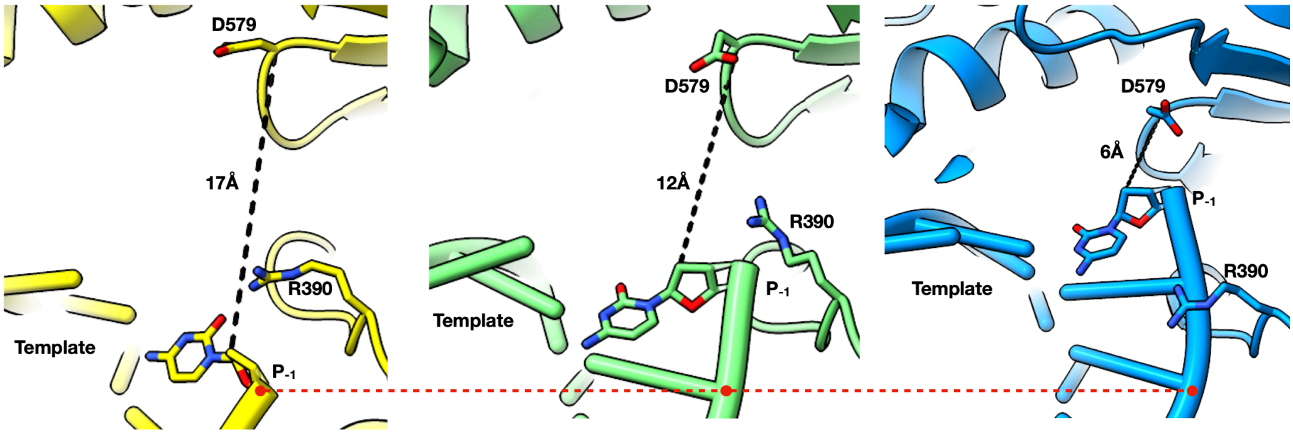


**Figure S4:** Distance between the 3' end of the primer strand and D579 (both shown in stick representation) in ternary1 (yellow; left), ternary2 (green; middle) and ternary3 (blue; right). For clarity the incoming dGTP has not been shown. Red dashed line indicates the position of the 3' end of the primer in ternary1. Position of R390 with respect to the primer strand is shown for reference.

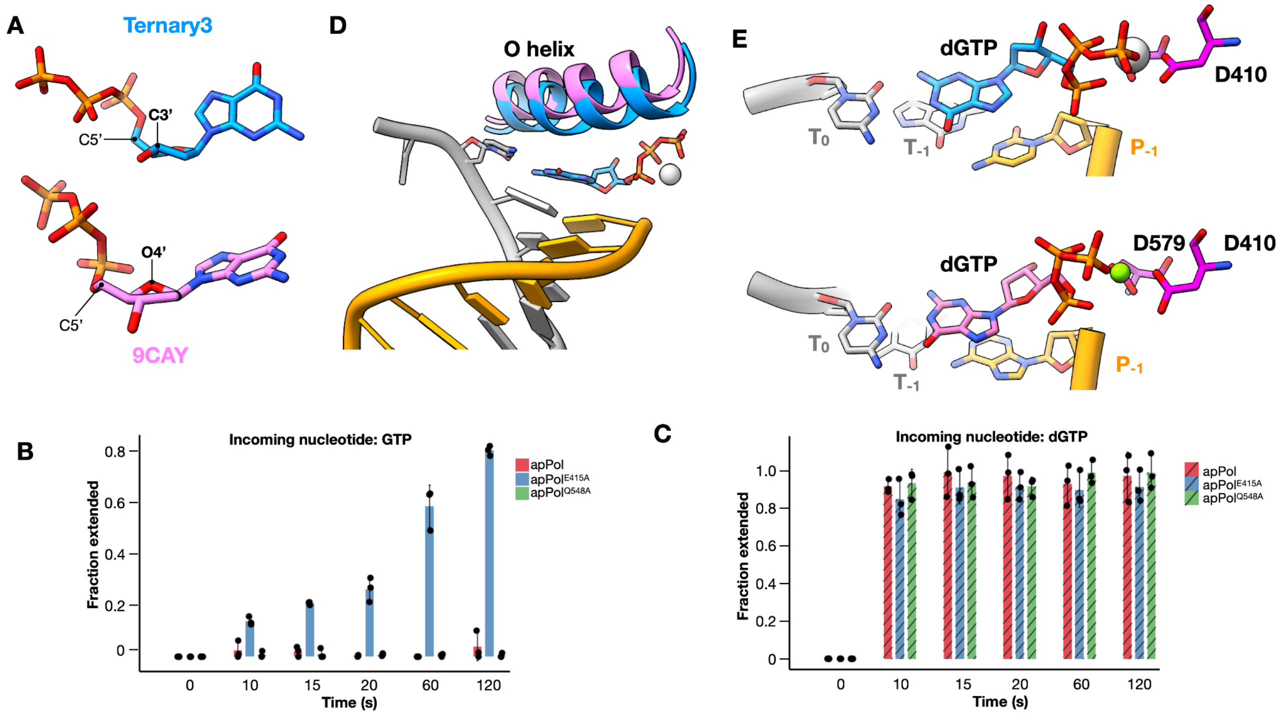


**Figure S5: Comparison between ternary3 and ternary4 (PDB id: 9CAY). (A)** Conformation of the incoming dGTP of ternary3 (top; blue) and 9CAY (bottom; pink). The C5' of the ribose is marked for reference. The carbon or oxygen atoms that are out of the plane of the furanose ring and pointing towards C5' (endo position) are highlighted in bold. **(B, C)** Time courses of GTP **(B)** or dGTP **(C)** incorporation by apPol (red), apPol$^{E415A}$ (blue) and apPol$^{Q548A}$ (green). 1µM enzyme was incubated with 50nM FAM-P/T and 125µM of dGTP or GTP was added to initiate the reaction. The reactions were quenched after various time intervals by adding 250 mM EDTA and the fraction of starting primer that has been extended by one nucleotide has been plotted as bars. The experiments were performed in triplicates and the individual replicates are shown as black circles on the respective bars (representing average of the three replicates). Error bars show the standard deviation. **(D)** Relative positions of the O helix in 9CAY (pink) and ternary3 (blue). The incoming dGTP (blue), $Ca^{2+}$ (white) and primer (orange)/template (grey) DNA of ternary3 are shown for reference. **(E)** Position of the incoming dGTP (blue for ternary3 and pink for 9CAY) with respect to the DNA substrate (primer: orange and template: grey) in 9CAY (top; ternary4) and ternary3 (bottom). The catalytic aspartates (magenta) and metal B (green for 9CAY and white for ternary3) are shown for reference. To maintain parity, the 3'OH of ternary3 primer strand has not been shown.

| Item | Citation |
|---|---|
| R studio | Posit team (2024). RStudio: Integrated Development Environment for R. Posit Software, PBC, Boston, MA. URL http://www.posit.co/. |
| ggplot2 | Wickham H (2016). *ggplot2: Elegant Graphics for Data Analysis*. Springer-Verlag New York. ISBN 978-3-319-24277-4, https://ggplot2.tidyverse.org. |
| dplyr | Wickham H, François R, Henry L, Müller K, Vaughan D (2023). *dplyr: A Grammar of Data Manipulation*. R package version 1.1.4, https://github.com/tidyverse/dplyr, https://dplyr.tidyverse.org. |
| readr | Wickham H, Hester J, Bryan J (2024). *readr: Read Rectangular Text Data*. R package version 2.1.5, https://github.com/tidyverse/readr, https://readr.tidyverse.org. |
| stringr | Wickham H (2023). *stringr: Simple, Consistent Wrappers for Common String Operations*. R package version 1.5.1, https://github.com/tidyverse/stringr, https://stringr.tidyverse.org. |
| ggpattern | FC M, Davis T, ggplot2 authors (2022). *ggpattern: 'ggplot2' Pattern Geoms*. https://github.com/coolbutuseless/ggpattern, https://coolbutuseless.github.io/package/ggpattern/index.html. |

**Table S1:** Software and R packages used for plotting primer extension data.

| | **Binary1** | **Binary2** | **Ternary1** | **Ternary2** | **Ternary3** | **Consensus** |
|---|---|---|---|---|---|---|
| **Data collection and processing** | | | | | | |
| Magnification | 81000X | 81000X | 81000X | 81000X | 81000X | 81000X |
| Voltage (kV) | 300 | 300 | 300 | 300 | 300 | 300 |

| | | | | | | |
|---|---|---|---|---|---|---|
| Electron exposure (e–/Å$^2$) | 70 | 70 | 70 | 70 | 60 | 70 |
| Defocus range (μm) | -0.8 to -3.8 | -0.8 to -3.8 | -0.8 to -3.8 | -0.8 to -3.8 | -1 to -3.8 | -0.8 to -3.8 |
| Pixel size (Å) | 1.06 | 1.06 | 1.06 | 1.06 | 1.06 | 1.06 |
| Symmetry imposed | None | None | None | None | None | None |
| Initial particle images (no.) | 1,922,870 | 1,922,870 | 1,922,870 | 1,922,870 | 1,993,221 | 1,922,870 |
| Final particle images (no.) | 53,452 | 20,205 | 9,780 | 10,241 | 92,787 | 354,534 |
| Map resolution (Å) | 3.7 | 3.9 | 4.2 | 4.2 | 3.5 | 3.2 |
| FSC threshold | | | | | | |
| Map resolution range (Å) | 2.8 to 5.0 | 3.0 to 5 | 3.5 to 6.0 | 3.5 to 6.0 | 2.7 to 6.0 | 2.5 to 5.0 |
| **Refinement** | | | | | | |
| Initial model used (PDB code) | 5DKT and 1LV5 | 5DKT | 5DKT and 1LV5 | 5DKT and 1LV5 | 5DKT and 1LV5 | 5DKT and 1LV5 |
| Model resolution (Å) | 4 | 4.2 | 4.4 | 4.6 | 4.1 | 3.4 |
| FSC threshold | 0.5 | 0.5 | 0.5 | 0.5 | 0.5 | 0.5 |
| Model resolution range (Å) | | | | | | |
| Map sharpening *B* factor (Å$^2$) | -133 | -114 | -124 | -134 | -155 | -169 |
| Model composition | | | | | | |
| Non-hydrogen atoms | 6022 | 5243 | 6072 | 6032 | 6006 | 5601 |
| Protein residues | 627 | 628 | 627 | 627 | 621 | 626 |
| Ligands | N/A | 1 | 1 | 1 | 2 | N/A |
| *B* factors (Å$^2$) | | | | | | |
| Protein | 72.04 | 100.6 | 109.66 | 120.11 | 78.18 | 80.89 |
| Nucleotide | 151.78 | N/A | 178.25 | 173.81 | 99.19 | 175.19 |
| Ligand | N/A | 114.94 | 124.95 | 135.29 | 76.81 | |
| R.m.s. deviations | | | | | | |
| Bond lengths (Å) | 0.005 | 0.004 | 0.004 | 0.004 | 0.005 | 0.004 |
| Bond angles (°) | 0.585 | 0.600 | 0.756 | 0.757 | 0.709 | 0.489 |
| Validation | | | | | | |
| MolProbity score | 1.42 | 1.43 | 1.47 | 1.70 | 1.64 | 1.39 |
| Clashscore | 4.42 | 4.17 | 4.9 | 6.45 | 5.54 | 5.58 |
| Poor rotamers (%) | 0.51 | 0.17 | 0.17 | 1.02 | 0.00 | 0.51 |
| Ramachandran plot | | | | | | |
| Favored (%) | 96.8 | 96.49 | 96.64 | 95.04 | 95.12 | 97.60 |
| Allowed (%) | 3.04 | 3.35 | 3.2 | 4.8 | 4.72 | 2.24 |
| Disallowed (%) | 0.16 | 0.16 | 0.16 | 0.16 | 0.16 | 0.16 |

**Table S2:** CryoEM map and atomic model refinement statistics.

**Movie 1: Overall domain motion of apPol as it traverses the pre-chemistry steps.** A morph between binary1, ternary1, ternary2 and ternary3 and back. The protein is in ribbon representation with the residues contacting the DNA strands in stick representation. In addition Y481, 485 and 486 are shown in stick form. Colour coding is the same as 1B with dGTP shown in dark green and the $P_{-3}$ position on the primer strand is highlighted in yellow.

**Movie 2: Motion of the thumb as apPol traverses the pre-chemistry steps.** A morph between binary1, ternary1, ternary2 and ternary3 and back, with a zoomed-in view of the thumb. Colour coding and residue representation are the same as Movie 1. Polar contacts between apPol and DNA are shown with black dashed lines.

**Movie 3: Corkscrew motion of the DNA as apPol traverses the pre-chemistry steps.** A morph between binary1, ternary1, ternary2 and ternary3 and back, with a zoomed-in view of the DNA duplex. The thumb domain has been omitted for clarity. Colour coding is the same as Movie 1. R390, D579 and D410 are shown in stick representation with colouring based on elements (oxygen: red, nitrogen: blue).

**Movie 4**: **Formation and dissolution of the pre-insertion checkpoint.** A morph between binary1, ternary1, ternary2 and ternary3 and back, with a zoomed-in view of the O1 helix. Colour coding and residue representation are the same as Movie 3. Polar contacts between apPol, dGTP and DNA are shown with black dashed lines.

**Movie 5**: **Interaction of the template strand with the palm domain.** A morph between binary1, ternary1, ternary2 and ternary3 and back, focusing on the interaction of the DNA template with the palm domain. Colour coding and residue representation are the same as Movie 1. Interactions between apPol, dGTP and DNA are shown with black dashed lines.

**Movie 6**: **DNA slippage with respect to the thumb.** A morph between binary1, ternary1, ternary2 and ternary3 and back, zooming in on the interactions of the DNA with the thumb helices H1, H2 and the loop connecting these two helices. Colour coding and residue representation are the same as Movie 1 with the following addition. Thumb residues interacting with the primer strand are colored based on the elements. Interactions between apPol, dGTP and DNA are shown with black dashed lines.
